# Incomplete Dominance of Deleterious Alleles Contributes Substantially to Trait Variation and Heterosis in Maize

**DOI:** 10.1101/086132

**Authors:** Jinliang Yang, Sofiane Mezmouk, Andy Baumgarten, Edward S. Buckler, Katherine E. Guill, Michael D. McMullen, Rita H. Mumm, Jeffrey Ross-Ibarra

## Abstract

**Abstract:** Deleterious alleles have long been proposed to play an important role in patterning phenotypic variation and are central to commonly held ideas explaining the hybrid vigor observed in the offspring by crossing two inbred parents. We test these ideas using evolutionary measures of sequence conservation to ask whether incorporating information about putatively deleterious alleles can inform genomic selection (GS) models and improve phenotypic prediction. We measured a number of agronomic traits in both the inbred parents and hybrids of an elite maize partial diallel population and re-sequenced the parents of the population. Inbred elite maize lines vary for more than 350,000 putatively deleterious sites, but show a lower burden of such sites than a comparable set of traditional landraces. Our modeling reveals widespread evidence for incomplete dominance at these loci, and supports theoretical models that more damaging variants are usually more recessive. We identify haplotype blocks using an identity-by-decent (IBD) analysis and perform genomic prediction analyses in which we weigh blocks on the basis of segregating putatively deleterious variants. Cross-validation results show that incorporating sequence conservation in genomic selection improves prediction accuracy for grain yield and other fitness-related traits as well as heterosis for those traits. Our results provide empirical support for an important role for incomplete dominance of deleterious alleles in explaining heterosis and demonstrate the utility of incorporating functional annotation in phenotypic prediction and plant breeding.

**Author Summary:** A key long-term goal of biology is understanding the genetic basis of phenotypic variation. Although most new mutations are likely disadvantageous, their prevalence and importance in explaining patterns of phenotypic variation is controversial and not well understood. In this study we combine whole genome-sequencing and field evaluation of a maize mapping population to investigate the contribution of deleterious mutations to phenotype. We show that *a priori* prediction of deleterious alleles correlates well with effect sizes for grain yield and that variants predicted to be more damaging are on average more recessive. We develop a simple model allowing for variation in the heterozygous effects of deleterious mutations and demonstrate its improved ability to predict both phenotypes and hybrid vigor. Our results help reconcile alternative explanations for hybrid vigor and highlight the use of leveraging evolutionary history to facilitate breeding for crop improvement.

## Introduction

Understanding the genetic basis of phenotypic variation is critical to many biological endeavors from human health to conservation and agriculture. Although most new mutations are likely deleterious [19], their importance in patterning phenotypic variation is controversial and not well understood [48]. Empirical work suggests that, although the long-term burden of deleterious variants is relatively insensitive to demography [61], population bottlenecks and expansion may lead to an increased abundance of deleterious alleles over shorter time scales such as those associated with domestication [1], postglacial colonization [36] or recent human migration [53]. Even when the impacts on total load are minimal, demographic change may have important consequences for the contribution of deleterious variants to phenotypic variation [41, 58, 61, 62]. Together, these considerations point to a potentially important role for deleterious variants in determining patterns of phenotypic variation, especially for traits closely related to fitness.

Maize (*Zea mays*) is an ideal system in which to study the impacts of deleterious variants. In addition to its global agricultural importance, maize has long been an important genetic model system [49] and central to debates about the basis of hybrid vigor and the role of deleterious alleles [2, 11]. The maize domestication bottleneck has lead to an increased burden of deleterious alleles in maize compared to its wild ancestor teosinte [67], and rapid expansion following domestication likely lead to an increase in new mutations and stronger purifying selection [1]. More recently, modern maize breeding has lead to dramatic reductions in effective population size [63], but inbreeding during the development of modern inbred lines may have decreased load by purging recessive deleterious alleles [8]. Nonetheless, substantial evidence suggests an abundance of deleterious alleles present in modern germplasm, from changes in heterozygosity during the process of inbreeding [25, 45] and selection [23] to genome-wide association results that reveal an excess of associations with genes segregating for damaging protein-coding variants [46].

Modern maize agriculture takes advantage of hybrid maize plants that result from the cross between two parental inbred lines [11]. These crosses result in a phenomenon known as hybrid vigor or heterosis, in which the hybrid plant shows improved agronomic qualities compared to its parents. Heterosis cannot be easily predicted from parental phenotype alone, and the genetic underpinnings of heterosis remain largely unknown. The most straightforward explanation for heterosis has been simple complementation of recessive deleterious alleles homozygous in one of the inbred parents [7, 12]. While this model is supported by considerable empirical evidence [22, 69], it fails in its simplest form to explain a number of observations, especially relating to heterosis and inbreeding depression in polyploid plants [2, 3, 70]. Other explanations, such as single-gene heterozygote advantage, clearly may play an important role in some cases [e.g. 31, 35], but mapping studies suggest such models are not easily generalizable [37].

In this study, we set out to investigate the contribution of deleterious alleles to phenotypic variation and hybrid vigor in maize. We created a partial diallel population from 12 maize inbred lines which together represent much of the ancestry of present-day commercial U.S. corn hybrids [43, 47]. We measured a number of agronomically relevant phenotypes in both parents and hybrids, including flowering time (days to 50% pollen shed, DTP; days to 50% silking, DTS; anthesis-silking interval, ASI), plant size (plant height, PHT; height of primary ear, EHT), grain quality (test weight which is a measure of grain density, TW), and grain yield (GY). We conducted whole genome sequencing of the parental lines and characterized genome-wide deleterious variants using genomic evolutionary rate profiling (GERP) [10]. We then test models of additivity and dominance for each phenotype using putatively deleterious variants and investigate the relationship between dominance and phenotypic effect size and the long-term fitness consequences of a mutation as measured by GERP. Finally, we take advantage of a Bayesian genomic selection framework [28] approach to explicitly test the utility of including GERP scores in phenotypic prediction for hybrid traits and heterosis.

## Materials and Methods

### Plant materials and phenotypic data

We formed a partial diallel population from the F1 progeny of 12 inbred maize lines (**Table S1**, Figure S1). Field performance of the 66 F1 hybrids and 12 inbred parents were evaluated along with two current commercial check hybrids in Urbana, IL over three years (2009-2011) in a resolvable incomplete block design with three replicates. To avoid competition effects, inbreds and hybrids were grown in different blocks within the field. Plots consisted of four rows (5.3 m long with row spacing of 0.76 m at plant density of 74,000 plants *ha*^−1^), with all observations taken from the inside two rows to minimize effects of shading and maturity differences from adjacent plots. We measured plant height (PHT, in cm), height of primary ear (EHT, in cm), days to 50% pollen shed (DTP), days to 50% silking (DTS), anthesis-silking interval (ASI, in days), grain yield adjusted to 15.5% moisture (GY, in bu/A), and test weight (TW, weight of 1 bushel of grain in pounds).

We estimated Best Linear Unbiased Estimates (BLUEs) of the genetic effects in ASReml-R (VSN International) with the following linear mixed model:

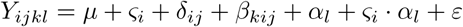

where *Y_ijkl_* is the phenotypic value of the *l^th^* genotype evaluated in the *k^th^* block of the *j^th^* replicate within the *i^th^* year; *μ*, the overall mean; *ς_i_*, the fixed effect of the *i^th^* year; *δ_ij_*, the random effect of the *j^th^* replicate nested within the *i^th^* year; *β_kij_*, the random effect of the *k^th^* block nested within the *i^th^* year and *j^th^* replicate; *α_l_*, the fixed genetic effect of the *l^th^* individual; *ς_i_* · *α_l_*, the random interaction effect of the *l^th^* individual with the *i^th^* year; and *ε*, the model residuals. We calculated the broad sense heritability (*H*^2^) of traits based on the analysis of all individuals (inbred parents, hybrid progeny, and checks) following the equation:

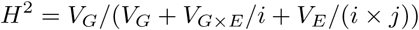

where *i* = 3 (number of years) and *j* = 3 (number of replicates per year).

The BLUE values for each cross can be found in **Table S1**; values across all hybrids were relatively normally distributed for all traits (Shapiro-Wilk normality tests *P* values > 0.05, Figure S1), though some traits were highly correlated (e.g. Spearman correlation *r* = 0.98 for DTS and DTP, Figure S2).

We estimated mid-parent heterosis (MPH) as:

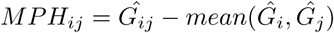

where 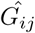, 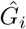 and 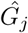 are the BLUE values of the hybrid and its two parents *i* and *j*. Note that for ASI, lower trait values are considered superior. General combining ability (GCA) was estimated following Falconer and Mackay [20], and the estimated values can be found in **Table S2**.

### Sequencing and Genotyping

We extracted DNA from the 12 inbred lines following [16] and sheared the DNA on a Covaris (Woburn, Massachusetts) for library preparation. Libraries were prepared using an Illumina paired-end protocol with 180 bp fragments and sequenced using 100 bp paired-end reads on a HiSeq 2000. Raw sequencing data are available at NCBI SRA (PRJNA381642).

We trimmed raw sequence reads for adapter contamination with Scythe (https://github.com/vsbuffalo/scythe) and for quality and sequence length (≥ 20 nucleotides) with Sickle (https://github.com/najoshi/sickle). We mapped filtered reads to the maize B73 reference genome (AGPv2) with bwa-mem [38], keeping reads with mapping quality higher than 10 and with a best alignment score higher than the second best one for further analyses.

We called single nucleotide polymorphisms (SNPs) using the *mpileup* function from samtools [39]. To deal with known issues with paralogy in maize [8], SNPs were filtered to be heterozygous in fewer than 3 inbred lines, have a mean minor allele depth of at least 4, have a mean depth across all individuals less than 30 and have missing alleles in fewer than 6 inbred lines. Data on total number of SNPs called and the rate of missing data per line are shown in **Table S3**. We estimated the allelic error rate using three independent data sets: for all individuals using 41,292 overlapping SNPs from the maize SNP50k bead chip [63]; for all individuals using 180,313 overlapping SNPs identified through genotyping-by-sequencing (GBS) [57]; and for B73 and Mo17 using 10,426,715 SNP from the HapMap2 project [8]. Alignments and genotypes for each of the 12 inbreds are available at CyVerse (https://de.cyverse.org/de/?type=data&folder=/iplant/home/yangjl/pvp_diallel_data). Because these parents are highly inbred, knowing their homozygous genotype also allows us to know the genotype of the F1 derived from any two of the parents.

To test whether alignment to the B73 reference introduces a bias in relatedness estimation, we computed kinship matrices using both our SNP data as well as genotyping-by-sequencing data (version AllZeaGBSv2.7 downloaded from (www.panzea.org)) obtained from alignments to a set of sequencing reads ascertained from a broad germplasm base [24]. The two matrices were nearly identical (Pearson’s correlation coefficient *r* = 0.995), suggesting the degree of relatedness among lines is not sensitive to using B73 as the reference genome.

### Identifying putatively deleterious alleles

We used genomic evolutionary rate profiling (GERP) [14] estimated from a multi-species whole-genome alignment of 13 plant genomes [55] including *Zea mays*, *Coelorachis tuberculosa*, *Vossia cuspidata*, *Sorghum bicolor*, *Oryza sativa*, *Setaria italica*, *Brachypodium distachyon*, *Hordeum vulgare*, *Musa acuminata*, *Populus trichocarpa*, *Vitis vinifera*, *Arabidopsis thaliana*, and *Panicum virgatum*; the alignment and estimated GERP scores are available at CyVerse (https://de.cyverse.org/de/?type=data&folder=/iplant/home/yangjl/pvp_diallel_data). We define “GERP-SNPs” as the subset of SNPs with GERP score > 0, and at each SNP we assign the minor allele in the multi-species alignment as the likely deleterious allele. Finally, we predicted the functional consequences of GERP-SNPs based on genome annotation information obtained from SnpEff [9]. The multi-species alignment made use of the B73 AGPv3 assembly, and to ensure consistent coordinates, we ported our SNP coordinates from AGPv2 to AGPv3 using the Gramene assembly converter (http://ensembl.gramene.org/Zea_mays/Tools/AssemblyConverter?db=core).

To compare GERP scores (for all SNPs with GERP > 0) to recombination rate and allele frequencies, we obtained the NAM genetic map [51] from the Panzea website (http://www.panzea.org/) and allele frequencies from the > 1,200 maize lines sequenced as part of HapMap3.2 [6]. To compare the burden of deleterious alleles in modern inbred lines to landraces, we extracted genotypic data of 23 specially-inbred traditional landrace cultivars (see [8] for more details) from HapMap3.2. For each line, we calculated burden as the count of minor alleles present across all GERP-SNPs divided by the total number of non-missing sites. We separated sites into fixed (present in all individuals of a group) and segregating sites for landrace and modern maize samples separately.

### Estimating effect sizes and dominance of GERP-SNPs

We estimated the additive and dominant effects of individual GERP-SNPs using a GBLUP model [13] implemented in GVCBLUP [65]:

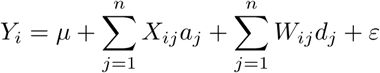

where *Y_i_* is the BLUE value of the *i*th hybrid, *a_j_* and *d_j_* are the additive and dominant effects of the *j*th GERP-SNP, *X_ij_* = {2*p*, 2*p* − 1, 2*p* − 2}, and *W_ij_* = {2*p*^2^,2*p*(1 − *p*),−2(1 − *p*)^2^} are the genotype encodings for genotypes *A*_1_*A*_1_, *A*_1_*A*_2_, and *A*_2_*A*_2_ of the *j*th SNP in the *i*th hybrid, respectively, and *ε* is the model residuals. The additive and dominance SNP encoding ensures that the effects are independent for a given GERP-SNP. We first estimated the total variance explained under models of complete additivity (*d_j_* = 0) or complete dominance (*a_j_* = 0). Then, to assess correlations between SNP effects and GERP scores, we calculated the degree of dominance (*k* = *d/a*) [42] for SNPs that each explained greater than the genome-wide mean per-SNP variance (total variance explained divided by total number of GERP-SNPs). Because this approach can lead to very large absolute values of *k*, we truncated GERP-SNPs with |*k* = *d/a*| > 2 for all further analyses.

To compare the variance explained by our model to that explained by random SNPs, we used a 2-dimensional sampling approach to create 10 equal-sized datasets of randomly sampled SNPs (including SNPs with GERP score <= 0) matched for allele frequency (in bins of 10%) and recombination rate (in quartiles of cM/Mb). For each dataset we fit the above model separately and estimated SNP effects and phenotypic variance explained by each SNP.

To test the relationship between GERP score and dominance under a simple model of mutation-selection equilibrium, we estimated the selection coefficient *s* by assuming that yield is a measure of fitness. We assigned the yield-increasing allele at each GERP-SNP a random dominance value in the range of 0 ≥ *k* ≥ 1 and calculated its equilibrium allele frequency *p* under mutation-selection balance using 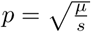 for values of *k* > 0.98 and 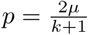 for *k* ≤ 0.98. We then simulated datasets using binomial sampling to choose SNPs in a sample of size *n* = 12 inbreds.

### Haplotype Analysis

We imputed missing data and identified regions of identity by descent (IBD) between the 12 inbred lines using the fastIBD method implemented in BEAGLE [4]. We then defined haplotype blocks as contiguous regions within which there were no IBD break points across all pairwise comparisons of the parental lines (Figure S3). Haplotype blocks at least 1 Kb in size were kept for further analyses.

Because there is no recombination in an inbred parent, this allows us to project the diploid genotype of each F1 based on the haplotypes of the two parents. In the projected diploid genotype of each F1, haplotype blocks were weighted by the summed GERP scores of all GERP-SNPs (python script ‘gerpIBD.py’ available at https://github.com/yangjl/zmSNPtools); blocks with no SNPs with positive GERP scores were excluded from further analysis. For a particular SNP with a GERP score *g*, the homozygote for the conserved (major) allele was assigned a value of 0, the homozygote for the putatively deleterious allele a value of 2*g*, and the heterozygote a value of (1 + *k*) × *g*, where *k* is the dominance estimated from the GBLUP model above.

### Genomic Selection

We used the BayesC option from GenSel4 [28] for genomic selection model training with 41,000 iterations and removing the first 1,000 as burn-in. We used the model

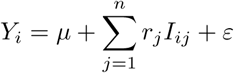

where *Y_i_* is the BLUE value of the *i*th hybrid, *r_j_* is the regression coefficient for the *j*th haplotype block, and *I_ij_* is the sum of GERP scores under an additive, dominance or incompletely dominance model for the *i*th hybrid in the *j*th haplotype block.

To conduct prediction, we used a 5-fold cross-validation method, dividing the diallel population randomly into training (80%) and validation sets (20%) 100 times. After model training, we obtained prediction accuracies by comparing the predicted breeding values with the observed phenotypes in the corresponding validation sets. For comparison, we permuted GERP scores using 50k SNP (≈ 100Mb or larger) windows which were circularly shuffled 10 times to estimate a null conservation score for each IBD block. We conducted permutations on all GERP-SNPs as well as on a restricted set of GERP-SNPs only in genic regions to control for GERP differences between genic (N = 221, 960) and intergenic regions (N = 123, 216). We conducted permutation cross-validation experiments using the same training and validation sets.

We estimated the posterior phenotypic variance explained using all of the data to derive correlations between breeding values estimated from the prediction model and observed BLUE values. Note that the correlation used here is different from the prediction accuracy (*r*) used for the cross-validation experiments, where the latter is defined as the correlation between real and estimated values; the two statistics will converge to the same value when there is no error in SNP/haplotype effect estimation [68].

Finally, to compare our genomic prediction model to a classical model of general combining ability, we used the following equations:

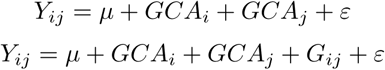

where *Y_ij_* is the BLUE value of the hybrid of the *i^th^* and *j^th^* inbreds, *μ* is the overall mean, *GCA_i_* and *GCA_j_* are the general combining abilities of the *i^th^* and *j^th^* inbreds, *G_ij_* is the breeding value of the hybrid of the *i^th^* and *j^th^* inbreds as estimated by our genomic prediction model, and *ε*, the model residuals.

### Data and code accessibility

Sequencing data have been deposited in NCBI SRA (SRP103329) database, and code for all analyses are available in the public GitHub repository (https://github.com/yangjl/GERP-diallel).

## Results

### Heterosis in a partial diallel population

We created a partial diallel population from 12 maize inbred lines which together represent much of the ancestry of present-day commercial U.S. corn hybrids (**Table S1**) [43, 47]. We measured a number of agronomically relevant phenotypes in both parents and hybrids, including flowering time (days to 50% pollen shed, DTP; days to 50% silking, DTS; anthesis-silking interval, ASI), plant size (plant height, PHT; height of primary ear, EHT), test weight (TW; a measure of quality based on grain density), and grain yield (GY). In an agronomic setting GY — a measure of seed production per unit area — is the primary trait selected by breeders and thus analogous to fitness. Plant height and ear height, both common measures of plant health or viability, were significantly correlated to GY (Figure S2).

For each genotype we derived best linear unbiased estimators (BLUEs) of its phenotype from mixed linear models (**Table S1**) to control for spatial and environmental variation (see Methods). We estimated mid-parent heterosis (MPH, Figure 1a) for each trait as the percent difference between the hybrid compared to the mean value of its two parents (see Methods, **Table S1**). Consistent with previous work [37], we find that grain yield (GY) showed the highest level of heterosis (MPH of 182% ± 60%). While flowering time (DTS and DTP) is an important adaptive phenotype globally [50], it showed relatively little heterosis in this study, likely due to the relatively narrow geographic range represented by the parental lines.

**Fig 1.**
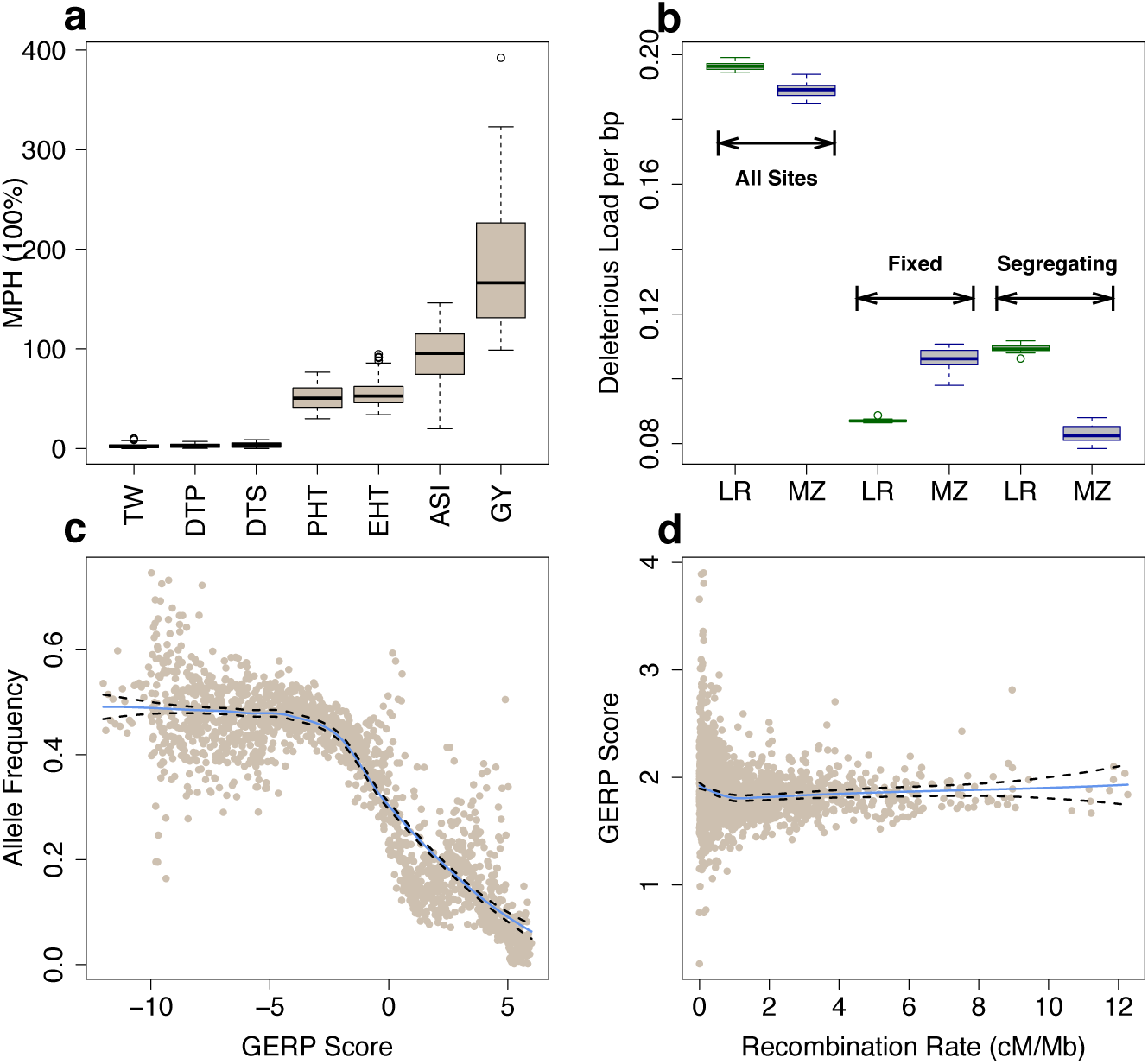
Heterosis and deleterious variants. (**a**) Boxplots (median and interquartile range) of percent mid-parent heterosis (MPH). (**b**) Proportion of deleterious alleles in landraces (LR, green) and elite maize (MZ, blue) lines. (**c**) The allele frequency of the minor alleles in the multi-species alignment in bins of 0.01 GERP score (including GERP < = 0 sites). (**d**) The mean GERP score for putatively deleterious sites (GERP > 0). Each point represents a 1 Mb window. In (**c**) and (**d**) the solid blue and dashed black lines define the best-fit regression line and its 95% confidence interval.

#### Annotation of deleterious alleles

We resequenced the 12 inbred parents to an average depth of ≈ 10×, resulting in a filtered set of 13.8M SNPs. Compared to corresponding SNPs identified by previous studies (see Methods), we observed a mean genotypic concordance rate of 99.1%. In order to quantify the deleterious consequences of variants *a priori*, we made use of Genomic Evolutionary Rate Profiling (GERP) [14] scores of the maize genome [56]. GERP scores provide a quantitative measure of the evolutionary conservation of a site across a phylogeny that allows characterization of the long-term fitness consequences of both coding and noncoding positions in the genome [34]. Sites with more positive GERP scores are inferred to be under stronger purifying selection, and SNPs observed at such sites are thus inferred to be more deleterious. At each site with GERP scores > 0 (hereafter called GERP-SNPs), we designated the minor allele from the multispecies alignment as putatively deleterious. Of the 350k total segregating GERP-SNPs in our parental lines, 14% are detected in coding regions, equally split between synonymous (N = 64,439) and non-synonymous (N = 65,376) sites (**Table S4**). Each line carries, on average, 139k potential deleterious SNPs (**Table S5**). The reference genome B73 contains only ≈ 1/3 of the deleterious SNPs of the other parents, likely due to reference bias in identifying deleterious variants. The F1 hybrids of the diallel each contain an average of ≈ 56, 000 homozygous deleterious SNPs, ranging from 47,219 (PH207 × PHG35) to 77,210 (PHG84 × PHZ51) (**Table S6**).

To compare the burden of deleterious variants between our elite maize lines and traditionally cultivated landraces, we used genotypes from the maize HapMap3.2 [6] for our diallel parents and 23 specially-inbred landrace lines [8] (**Table S5**). Compared to landraces, the parents of our diallel exhibited a greater burden of fixed (allele frequency of 1) deleterious variants but a much smaller burden of segregating SNPs, resulting in a slightly lower overall proportion of deleterious sites (mean of 1.3M deleterious alleles out of 6.5M total sites vs. 0.6/3.3M; Figure 1b).

Population genetic theory predicts that deleterious variants should be at low overall frequencies, and that such variants should be enriched in regions of the genome with extremely low recombination [29]. Using data from more than 1,200 lines in maize HapMap3.2 [6], we find that allele frequency of the minor alleles in the multi-species alignment shows a strong negative correlation with GERP score (Figure 1c). This negative correlation holds using allele frequency derived from our 12 parental lines (Figure S4), though as expected is less significant given the smaller sample size. SNPs found in regions of the genome with low recombination also show higher overall GERP scores (Figure 1d), a trend particularly noticeable around centromeres (Figure S5). These results match previous empirical findings in maize that deleterious alleles are rare [46] and most abundant in the lowest recombination regions [25, 44, 55], and support the use of GERP scores as a quantitative measure of the long-term fitness effects of an observed variant.

#### Phenotypic effects of deleterious SNPs

We first investigate the impacts of deleterious variants on phenotype using simple linear regressions. Across all hybrids, the number of homozygote GERP-SNPs was negatively correlated with grain yield, plant height, and ear-height *per se* (see **Table S6** for complementation data and **Table S7** for correlations with all traits).

We next applied a genomic best linear unbiased prediction (GBLUP) [13] modeling approach to estimate the effect sizes and variance explained by GERP-SNPs for each of the phenotypes *per se* across our diallel (see Methods). GERP-SNPs had larger average effects and explained more phenotypic variance than the same number of randomly sampled SNPs (including SNPs with GERP score < = 0) matched for allele frequency and recombination (Figure 2a). We found the cumulative proportion of dominance variance explained by GERP-SNPs was higher for traits showing high heterosis (Spearman correlation *P* value < 0.01, *r* = 0.9), from ≈ 0 for flowering time traits to as much as 24% for grain yield (Figure S6). Distributions of per-SNP dominance 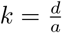 (see Methods) across traits were consistent with the cumulative partitioning of variance components (Figure 2b) and matched well with expectations from previous studies showing a predominantly additive basis for flowering time [5] and plant height [52] but meaningful contributions of dominance to test weight and grain yield [37, 43]. Although our diallel population is relatively small, our estimated values explain as much (for traits with low dominance variance like flowering time) or more variance (for traits with substantial dominance variance like grain yield) than sets of data with randomly shuffled values of dominance (n=10 randomizations of *k* per trait; Figure S7).

**Fig 2.**
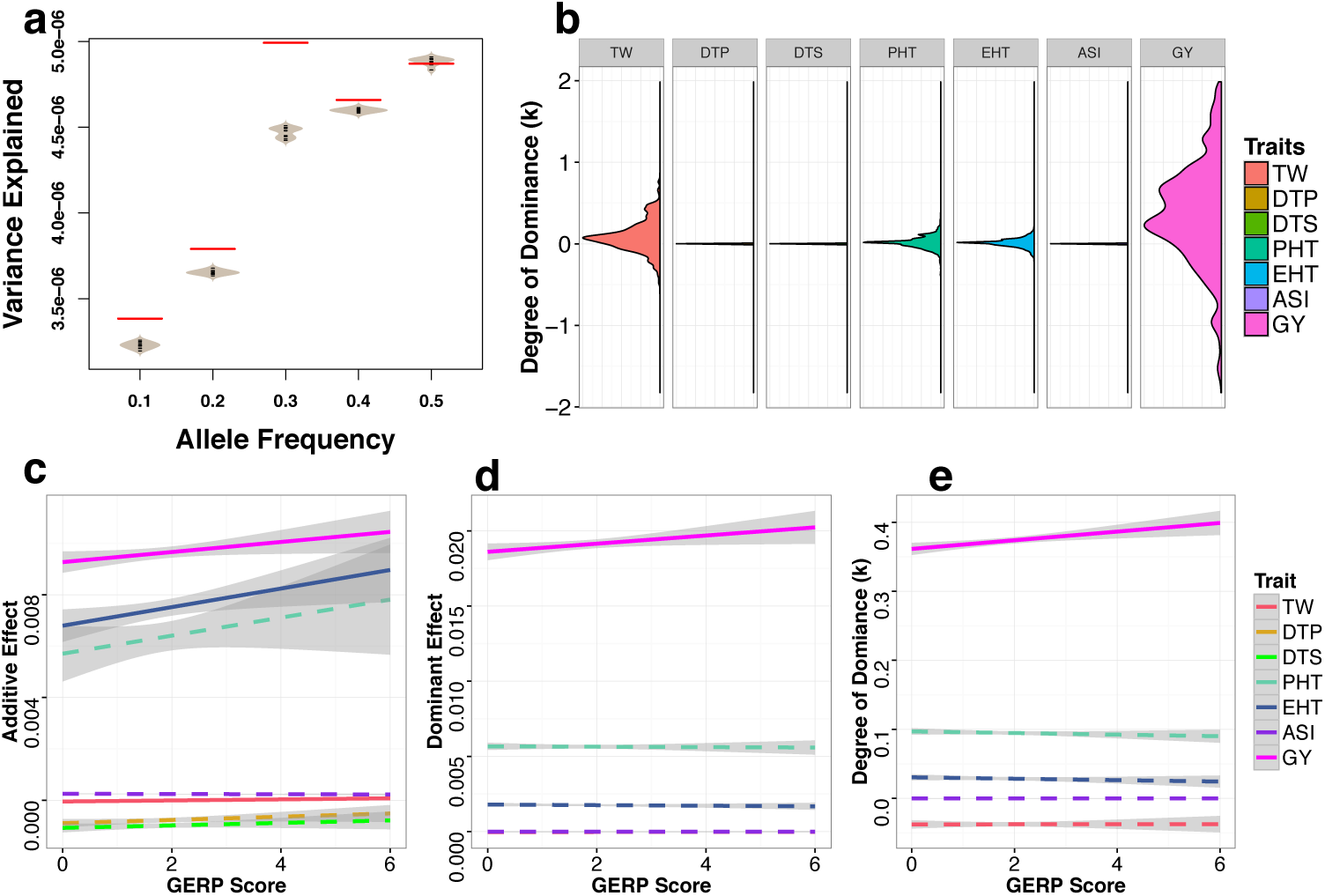
Variance explained and degree of dominance (*k*) of GERP-SNPs for traits *per se*. **(a)** Total per-SNP variance explained for grain yield trait *per se* by GERP-SNPs (red lines) and randomly sampled SNPs (grey beanplots). **(b)** Density plots of the degree of dominance (*k*). Extreme values of *k* were truncated at 2 and −2. **(c-e)** Linear regressions of additive effects **(c)**, dominance effects **(d)**, and degree of dominance **(e)** of seven traits *per se* against SNP GERP scores. Solid and dashed lines represent significant and nonsignificant linear regressions, with grey bands representing 95% confidence intervals. Data are only shown for SNPs that explain more than the mean genome-wide per-SNP variance (see Methods for details).

We then evaluated the relationship between GERP score and SNP effect size, dominance, and contribution to phenotypic variance. We found weak or negligible correlations between effect size and GERP score for flowering time and grain quality, but a strong positive correlation for fitness-related traits (Figure 2c-d). The variance explained by individual SNPs, however, was largely independent of GERP score (Figure S8), likely due to the observed negative correlation between allele frequency and GERP score (Figure 1c). Finally, we observed a positive relationship between GERP score and the degree of dominance (*k*) for grain yield (Figure 2e), such that the putatively deleterious allele at SNPs with higher GERP scores are also estimated to be more recessive for their phenotypic effects on grain yield (larger *k* for the major allele).

We investigated a number of possible caveats to the results presented in Figure 2. First, to control for the potential inflation of SNP effect sizes in regions of high linkage disequilibrium, we removed SNPs from regions of the genome in the lowest quartile of recombination. While some individual correlations changed significance, our overall results appear robust to the removal of low recombination regions (Figure S9). Second, we tested the impact of reference bias caused by inclusion of the B73 genome in the multi-species alignment used to estimate GERP scores. To do so, we removed the 11 hybrids which include as one parent the reference genome line B73 and repeated the above analyses. Doing so dramatically reduces the size of our dataset, but we nonetheless find significant correlations between complementation and phenotype (**Table S7**), that GERP-SNPs explain a greater proportion of overall variation than randomly sampled SNPs (Figure S10a), and that the relative pattern of dominance among traits remains the same (Figure S10b). While most of the correlations between effect size and GERP score lose significance (Figure S10c-d), likely due to the decreased sample size, the positive correlation between dominance and GERP score remains significant even in the absence of B73-derived hybrids (Figure S10e). Finally, because natural selection will maintain dominant deleterious alleles at lower frequencies than their recessive counterparts, we investigated whether the ascertainment bias against rare alleles present in our small sample would lead to the observed correlation between GERP and dominance. Simulations of SNPs with random dominance at mutation-selection balance (see Methods), however, failed to find any relationship between dominance and GERP score (Figure S11), though we caution that the dramatic demographic shifts involved in the recent history of maize [1] make such a simulation approximate at best.

#### Genomic prediction by incoporating GERP information

To explicitly test the informativeness of alleles identified *a priori* as putatively deleterious, we implemented a haplotype-based genomic prediction model that incorporates GERP scores as weights (see Methods). We explored the explanatory power with several different models and found that a model which incorporates both GERP scores and dominance (*k*) estimated from our GBLUP model explained a greater amount of the posterior phenotypic variance for most traits *per se* (Figure 3a) and heterosis (MPH) (Figure 3b). A simple additive model showed superior explanatory power for flowering time, however, consistent with previous association mapping results that flowering time traits are predominantly controlled by a large number of additive effect loci [5].

**Fig 3.**
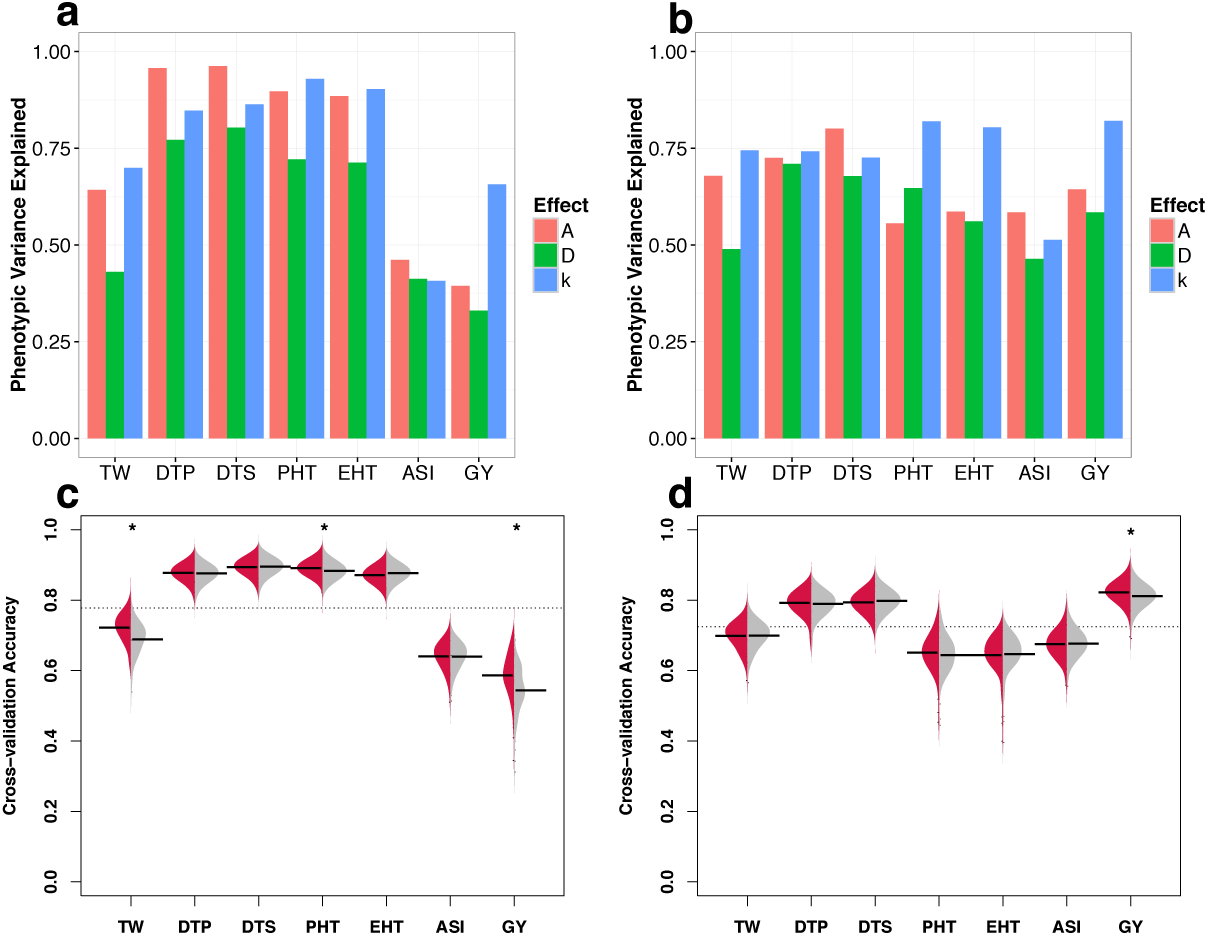
Genomic prediction models incorporating GERP. **(a-b)** Total phenotypic variance explained for traits *per se* **(a)** and heterosis (MPH) (**b**) under models of additivity (red), dominance (green), and incomplete dominance (blue). **(c-d)** Beanplots represent prediction accuracy estimated from cross-validation experiments for traits *per se* **(c)** and heterosis (MPH) **(d)** under a model of incomplete dominance. Prediction accuracy using estimated values for each GERP-SNP under an incomplete dominance model is shown on the left (red) and permutated values on the right (grey). Horizontal bars indicate mean accuracy for each trait and the grey dashed lines indicate the overall mean accuracy. Stars above the beans indicate prediction accuracies significantly (FDR < 0.05) higher than permutations. Results for pure additive and dominance models are shown in Figure S13.

To explicitly test the utility of incorporating GERP information in prediction models, we compared cross-validation prediction accuracies of the observed GERP scores to those from datasets in which GERP scores were circularly shuffled along the genome (see Methods). Models incorporating our observed GERP scores out-performed permutations (Figure 3c-d), even when considering only SNPs in genes (Figure S12). Our model improved prediction accuracy of grain yield by more than 4.3%, and improvements were also seen for plant height (0.8%) and testing weight (3.3%). While our model showed no improvement in predicting heterosis for traits showing low levels of heterosis (Figure 1a), including GERP scores significantly improved prediction accuracy for heterosis of grain yield (by 1%). Finally, our approach also significantly improved model fit for phenotypes of all traits *per se* as well as heterosis for GY and PHT compared to traditional models of genomic selection that use general combining ability (see Methods, **Table S2**) calculated directly from the pedigree of the hybrid population [26] (ANOVA FDR < 0.01 and difference in AIC < 0, **Table S8**).

## Discussion

We combine *a priori* prediction of deleterious alleles from whole genome sequence data with multi-year field evaluation of important agronomic phenotypes to test the role of incomplete dominance in determining hybrid phenotypes and heterosis in maize.

We first show that GERP scores are meaningful quantitative estimates of the fitness consequences of individual alleles, as SNPs with higher GERP scores are found at lower allele frequencies (Figure 1c), enriched in regions of low recombination (Figure 1d), and associated with larger effect sizes on grain yield (Figure 2c,d). Although a number of other methods exist to identify deleterious alleles from sequence data, GERP scores include both coding and noncoding sequence, do not require additional functional annotation, and show higher sensitivity and specificity than other related approaches [34]. While the GERP scores used here reflect conservation of across relatively deep phylogenetic time, future efforts may be able to increase power by incorporating information from within-species polymorphism data [32] as well as other types of annotations that have been shown to contribute substantially to phenotypic variation (e.g. Wallace *et al.*, [64] and Rodgers-Melnick *et al.*, [56]).

Using GERP scores as a proxy for deleterious alleles, we then ask whether our elite maize inbreds show an increased burden of deleterious alleles compared to a set of traditional landrace varieties. We find that modern inbreds are characterized by an increase in the proportion of deleterious variants fixed within the population (Figure 1b), consistent with the strong impact of drift associated with rapid decreases in effective population size during modern breeding [63]. In contrast, modern maize inbreds exhibit a much smaller proportion of segregating deleterious variants than landraces. This latter result is likely due to increased inbreeding in smaller populations, an effect exacerbated by the transition from traditional open-pollinated maize to the intentional formation of inbred lines. Inbreeding facilitates the removal of deleterious variants by selection, as evidenced by the striking inbreeding depression exhibited by open-pollinated maize [17]. Supporting our interpretation of these results, our observed differences in the burden of deleterious variants closely mimic results from simulations of partially recessive deleterious alleles in populations that have recently undergone demographic bottlenecks [61].

We next use the set of SNPs with GERP > 0 scores (or GERP-SNPs) to investigate the phenotypic effects of deleterious variants. Across phenotypes, our results largely mirror previous work, finding that dominance contributes substantially to grain yield [37], while traits such as flowering time appear to be largely additive [5]. At the level of individual SNPs we find correlations between GERP score and phenotypic effect size for yield and ear height, suggesting that long-term evolutionary constraint as measured by GERP is a useful predictor of the phenotypic effects of variants on traits related to fitness. Both traits are well explained by a model allowing for incomplete dominance (Figure 3a), as is plant height, which shows a positive but not significant correlation between effect size and GERP score. For grain yield, we also find that more deleterious alleles (those with higher GERP score) are more likely to be recessive. We are unaware of previous demonstrations of the genome-wide relationship between dominance and fitness in other multicellular organisms, but this result follows predictions based on models of metabolic pathways [33] and supports previous empirical evidence from gene knockouts in yeast [54]. Though our population size is small, our partial diallel crossing design and genomic selection model circumvent some of the problems with standard genome-wide association analyses, including genome-wide multiple testing thresholds and difficulties in assessing the effects of rare alleles due to limited replication. And while there is likely substantial error in individual SNP estimates, permutation analyses show our overall results nonetheless produce meaningful results (Figure 2a, Figure S7).

After showing that GERP-SNPs explain a substantial portion of the observed phenotypic variation when combined with our estimates of dominance and effect size, we more rigorously test the direct utility of GERP scores using cross-validation prediction methods. We show that for both plant height and grain yield, our GERP-enabled prediction model has significantly improved accuracy compared to randomized data, even when only considering SNPs within genes (which have higher on average GERP scores; Figure S12). As genotyping costs continue to decline, genomic prediction models are increasing in popularity [15]. Most previous work on genomic prediction, however, focuses exclusively on statistical properties of the models, ignoring potentially useful biological information (but see Edwards *et al.,* [18] for a recent example). Our results suggest that incorporating additional annotations — in particular information on evolutionary constraint — can provide additional, inexpensive benefits to existing genomic prediction frameworks.

Finally, our results also have implications for understanding the genetic basis of heterosis. Heterosis has been observed across many species, from yeast [59] to plants [60] and vertebrates [21], and a number of hypotheses have been put forth to explain the phenomenon [2, 7]. Of all these explanations, complementation of recessive deleterious alleles [7, 11] remains the simplest genetic explanation and is supported by considerable empirical evidence [22, 66, 69]. It remains controversial, however, because complementation of completely recessive mutations cannot fully explain a number of empirical observations including unexpected differences in heterosis and inbreeding depression among polyploids [2, 40]. For example, a model of simple complementation of purely recessive alleles is unable to explain differing levels of heterosis between triploid hybrids with different numbers of parental genomes (e.g. AAB vs ABB) [70] or why the cross of two tetraploid F1 hybrids shows greater heterosis than the original F1 [3]. Our results, however, indicate that most deleterious SNPs show incomplete dominance (Figure 2b) for traits with high levels of heterosis, and our genomic prediction models find improvement in predictions of heterosis when incorporating GERP scores under such a model (Figure 3d). These results are in line with other empirical evidence suggesting that new mutations tend to be partially recessive [30] and that GWAS hits exhibit incomplete dominance for phenotypes *per se* among hybrids [31]. We argue that allowing for incomplete dominance effectively unifies models of simple complementation with those of gene dosage [70]. Combined with observations that deleterious SNPs are enriched in low-recombination pericentromeric regions [55] (Figure 1d), such a model can satisfactorily explain changes in heterozygosity during breeding [23, 44], enrichment of yield QTL and apparent overdominance in centromeric regions [37], and even observed patterns of heterosis in polyploids (Figure S14). It is unlikely of course that any single explanation is sufficient for a phenomenon as complex as heterosis, and other processes such as overdominance likely make important contributions (e.g. Guo *et al.,* [27] and Huang *et al.,* [31]), but we argue here that a simple model of incompletely dominant deleterious alleles may provide substantial explanatory power not only for fitness-related phenotypic traits but for hybrid vigor as well.

## Conclusion

In this study, we use genomic and phenotypic data from a partial diallel population of maize to show that an incomplete dominance model of deleterious mutation both fits predictions of population genetic theory and explains phenotypic variation for fitness-related phenotypes and hybrid vigor. We find genome-wide support for hypotheses predicting that more damaging variants are more recessive. Finally, we show that leveraging evolutionary annotation information *in silico* enables us to predict grain yield and other traits, including heterosis, with greater accuracy. Together, these results help reconcile alternative explanations for hybrid vigor and point to the utility of leveraging evolutionary history to facilitate breeding for crop improvement.

## Acknowledgements

We would like to thank Tim Beissinger, Graham Coop, James Holland, Matthew Hufford, Emily Josephs, Peter Morrell, Michelle C. Stitzer, Kevin Thornton, and Stephen Wright for helpful discussion.

## 1 Supporting Tables

**Table S1.** BLUE values and levels of heterosis of the seven phenotypic traits for the 66 hybrids. (https://github.com/yangjl/GERP-diallel/blob/master/table/Table_trait_heterosis.csv)

**Table S2.** General combining ability and specific combining ability of the seven phenotypic traits. (https://github.com/yangjl/GERP-diallel/blob/master/table/Table_CA.csv)

**Table S3.** SNP missing rate in our diallel parental lines. (https://github.com/yangjl/GERP-diallel/blob/master/table/Table_SNP_missing_rate.csv)

**Table S4.** Summary statistics of SNP annotation results. (https://github.com/yangjl/GERP-diallel/blob/master/table/Table_S_snpeff_results.xlsx)

**Table S5.** Number of deleterious SNPs carried per line. (https://github.com/yangjl/GERP-diallel/blob/master/table/Table_S_del_per_line.csv)

**Table S6.** Number of complementation and homozygote deleterious load for GERP-SNPs in hybrids. (https://github.com/yangjl/GERP-diallel/blob/master/table/Table_S_del_complemenation.csv)

**Table S7.** The correlation between the number of homozygote GERP-SNPs and the hybrid phenotypes. (https://github.com/yangjl/GERP-diallel/blob/master/table/Table_hyb_load_pheno.csv)

**Table S8.** Model comparisons *P* values and AICs. (https://github.com/yangjl/GERP-diallel/blob/master/table/Table_model_comp.csv)

## 2 Supporting Figures

**Fig S1.**
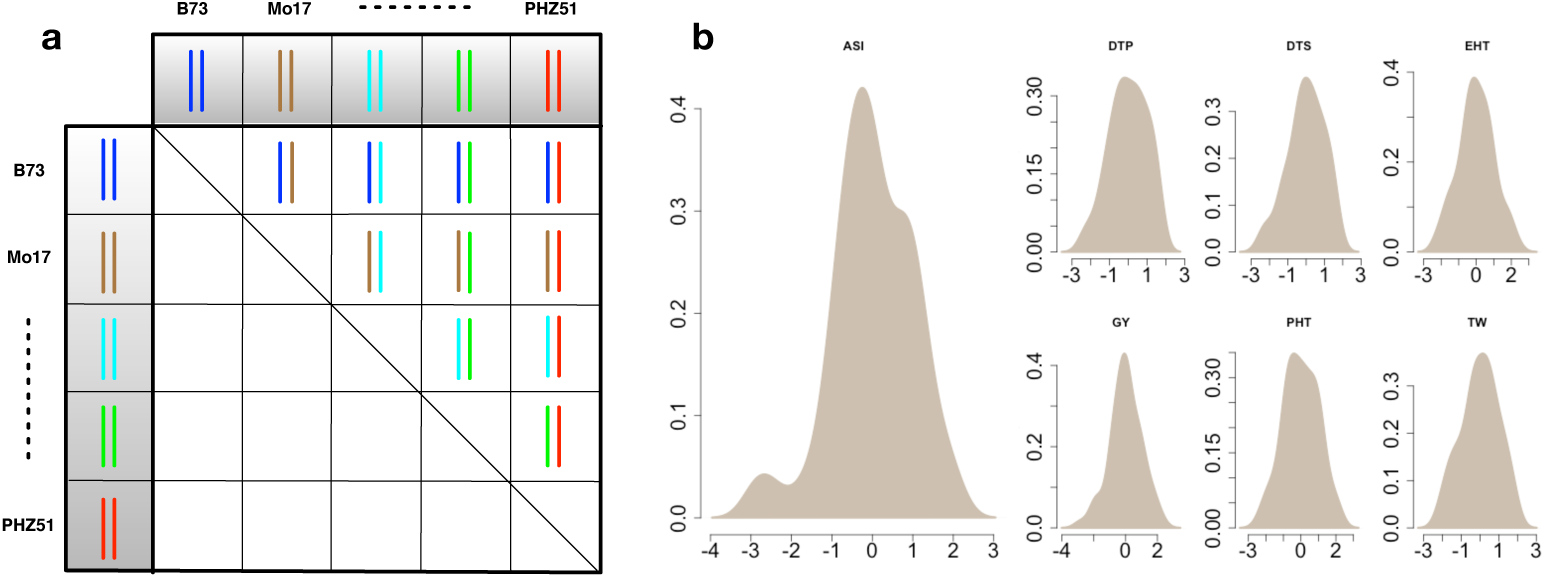
A half-diallel population and distributions of phenotypes. (**a**) Twelve maize inbred lines were selected and crossed in a half-diallel fashion. Each inbred lines was used as both male and female and the resulting F1 seed was bulked. (**b**) Density plots of normalized BLUE values for the seven phenotypic traits. We used “scale” function in R to normalize the BLUE values by first centering on zero and then dividing the numbers by their standard deviation. The seven phenotypic traits are plant height (PHT), height of primary ear (EHT), days to 50% pollen shed (DTP), days to 50% silking (DTS), anthesis-silking interval (ASI), grain yield adjusted to 15.5% moisture (GY), and test weight (TW).

**Fig S2.**
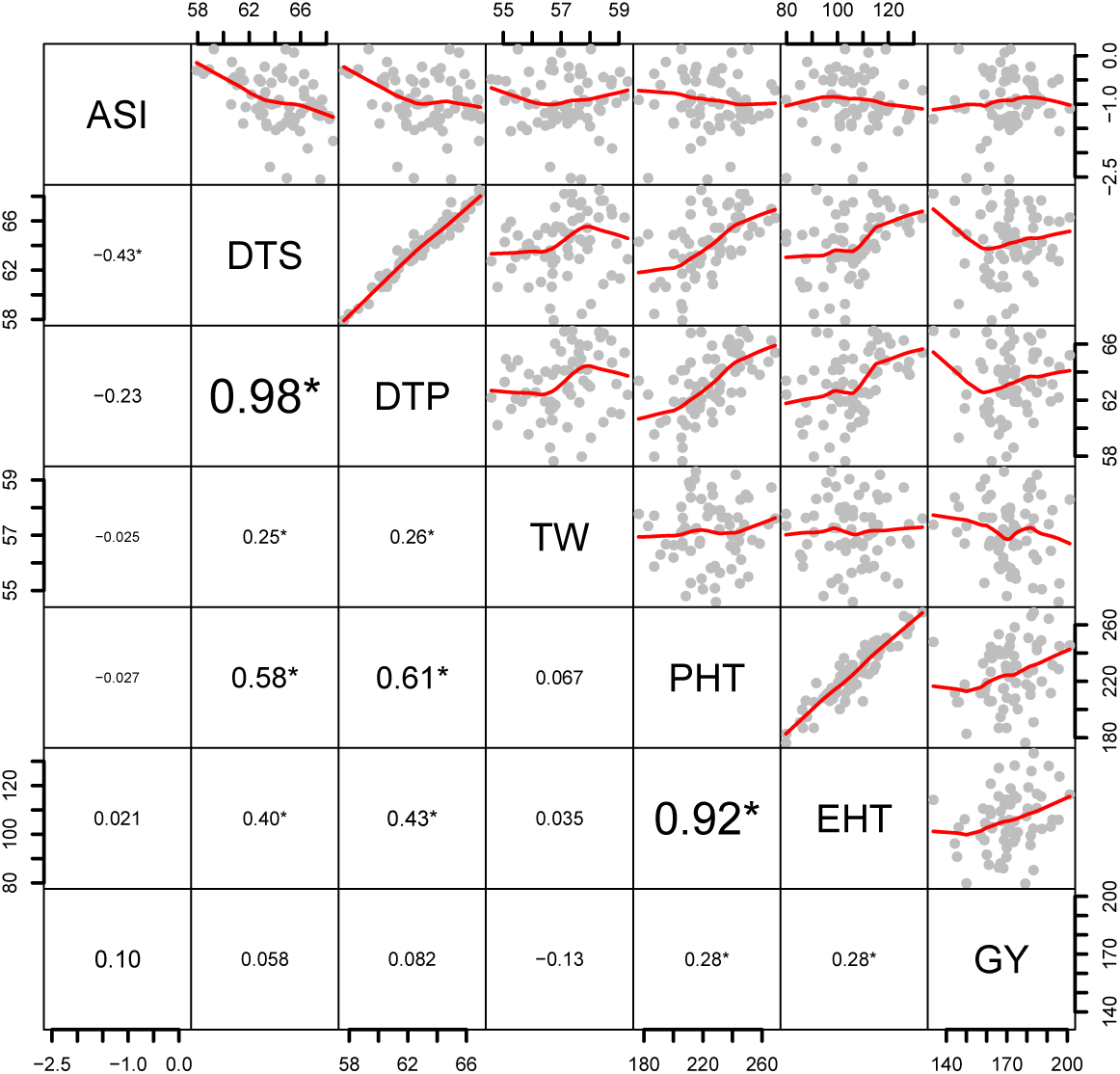
Pairwise correlation plots of seven phenotypic traits. The upper right panels show the scatter plots of all possible pairwise comparisons of two traits. Red line is a fitted smooth curve using “loess” method. In the lower left panels, the numbers are the Spearman correlation coefficients (*r*) and the asterisks (*) indicate the correlation coefficients are statistically significant (Spearman correlation test *P* value < 0.05). Units for various traits are plant height (PHT, in cm), height of primary ear (EHT, in cm), days to 50% pollen shed (DTP), days to 50% silking (DTS), anthesis-silking interval (ASI, in days), grain yield adjusted to 15.5% moisture (GY, in bu/A), and test weight (TW, weight of 1 bushel of grain in pounds).

**Fig S3.**
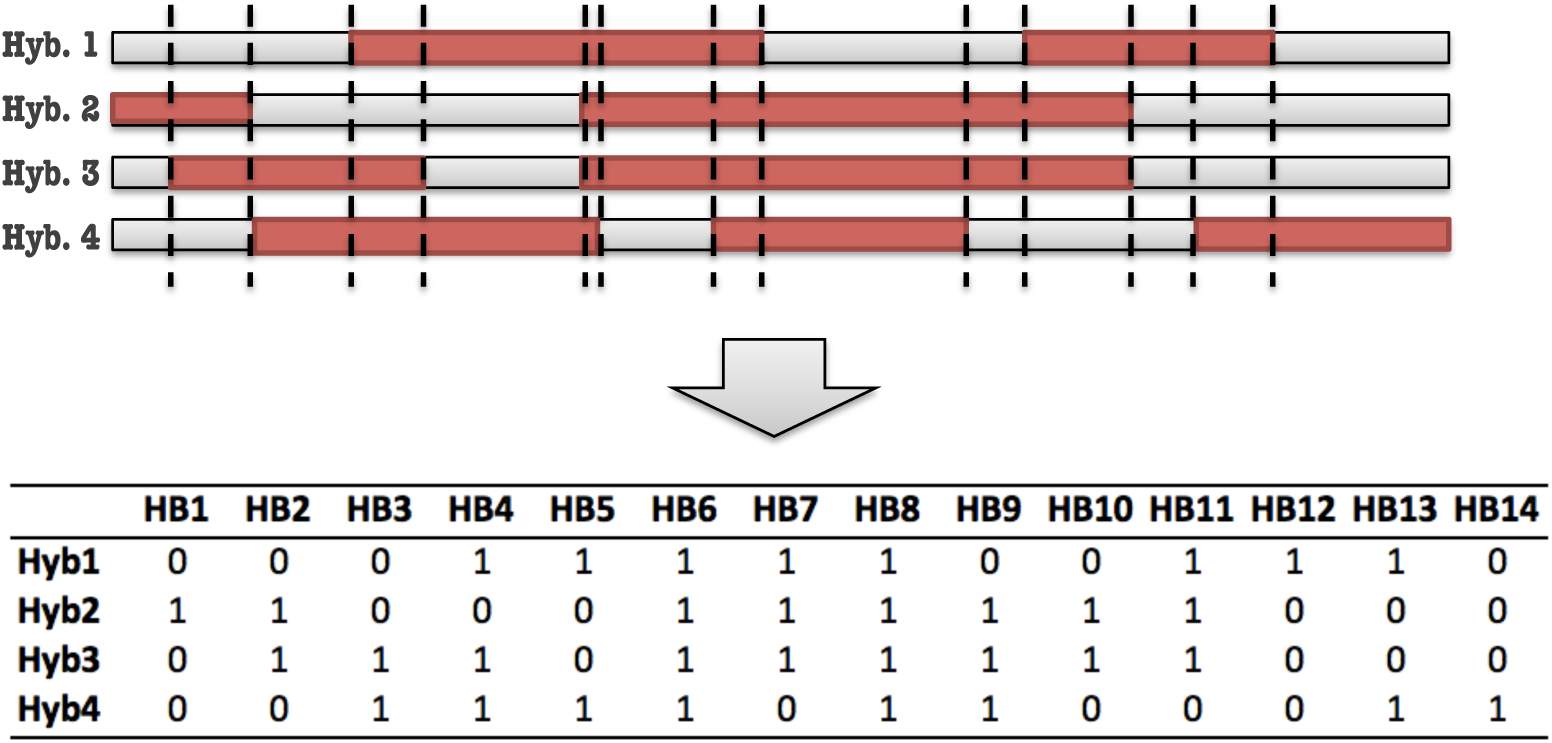
Haplotype block identification using an IBD approach. In the upper panel, regions in red are IBD blocks identified by pairwise comparison of the two parental lines of a hybrid. The vertical dashed lines define haplotype blocks. In the lower panel, hybrid genotype in each block are coded as heterozygotes (0) or homozygotes (1).

**Fig S4.**
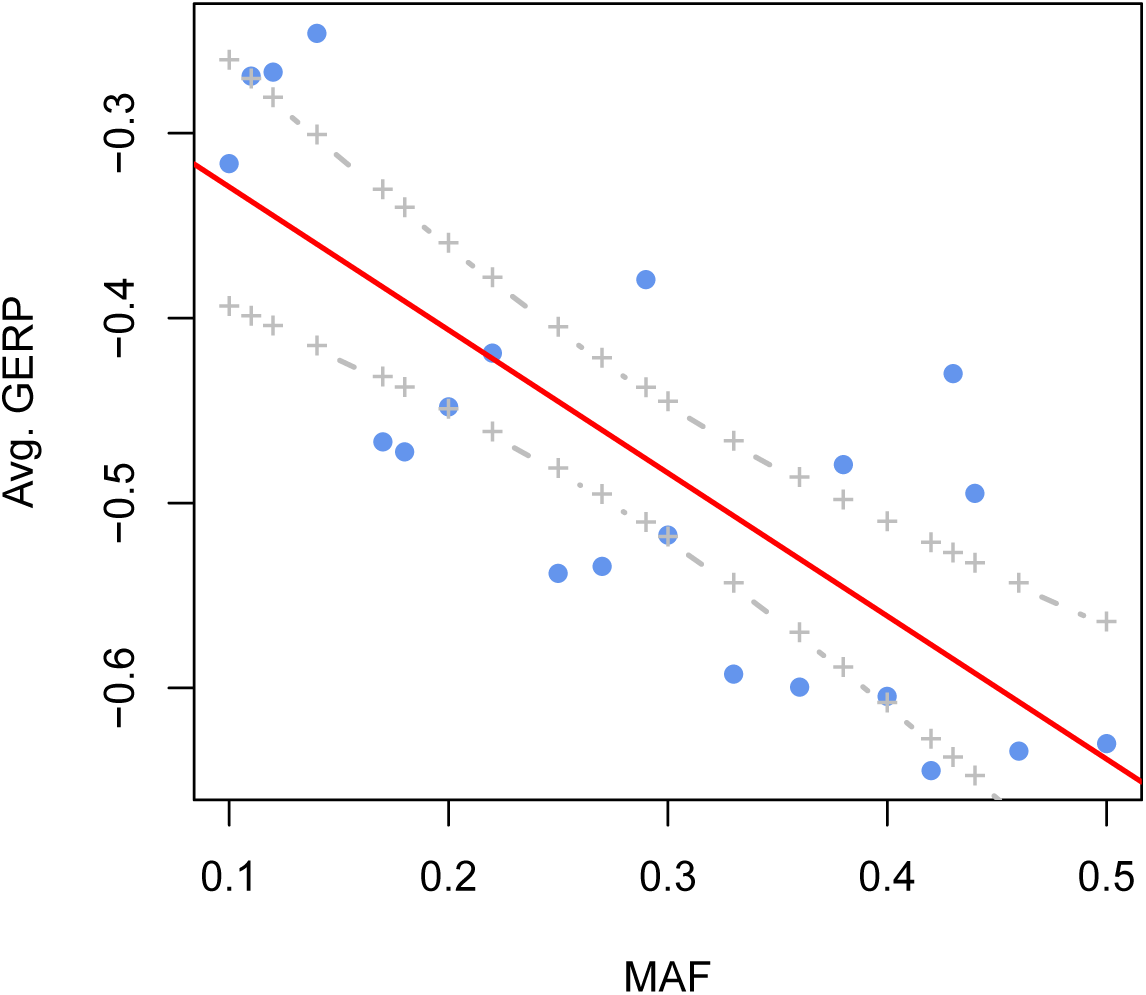
The minor allele frequency estimated from 12 parental lines in bins of 0.01 GERP score. Red solid and grey dashed lines define the best-fit regression line and its 95% confidence interval.

**Fig S5.**
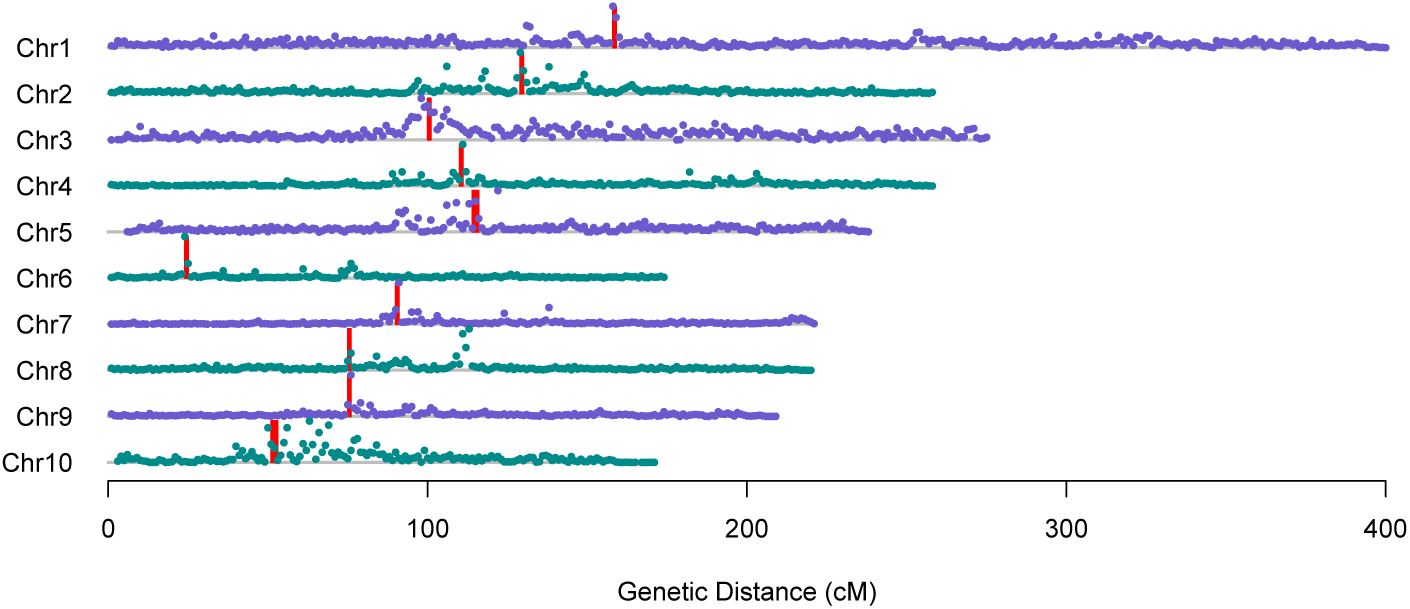
Segregating genetic load across ten maize chromosomes. Dots indicate mean GERP scores of putatively deleterious SNPs (GERP scores > 0) carried by our 12 elite maize lines (bin size = 1 cM). Vertical red lines indicate centromeres.

**Fig S6.**
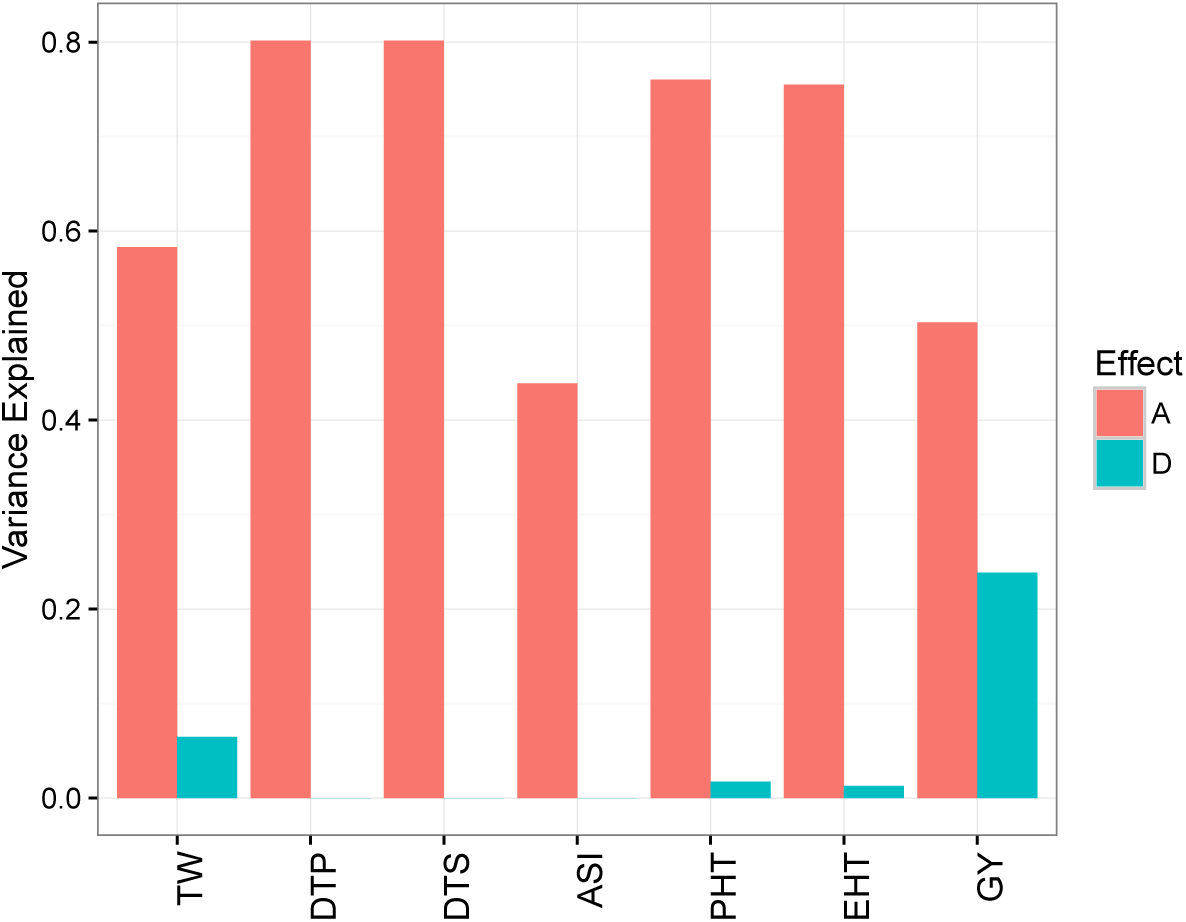
Cumulative variance explained by GERP-SNPs. Additive and dominance effects are indicated by red and blue colors respectively.

**Fig S7.**
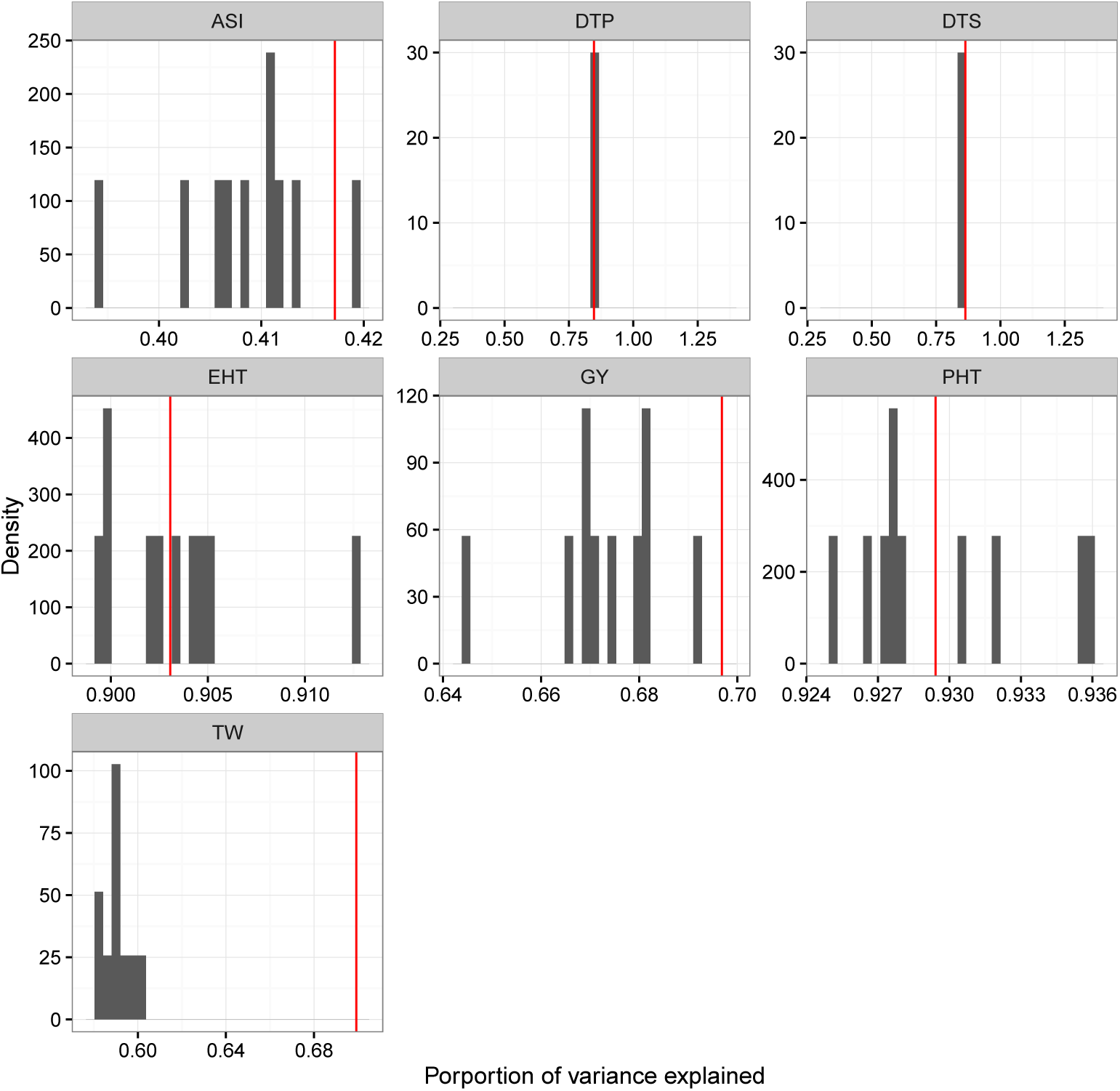
Phenotypic variance explained for observed data and for randomly shuffled data using the genomic selection model. Histograms show the results for the randomly shuffled (10 times) degrees of dominance (*k*) in each trait. Red lines are phenotypic variance explained using the observed *k*.

**Fig S8.**
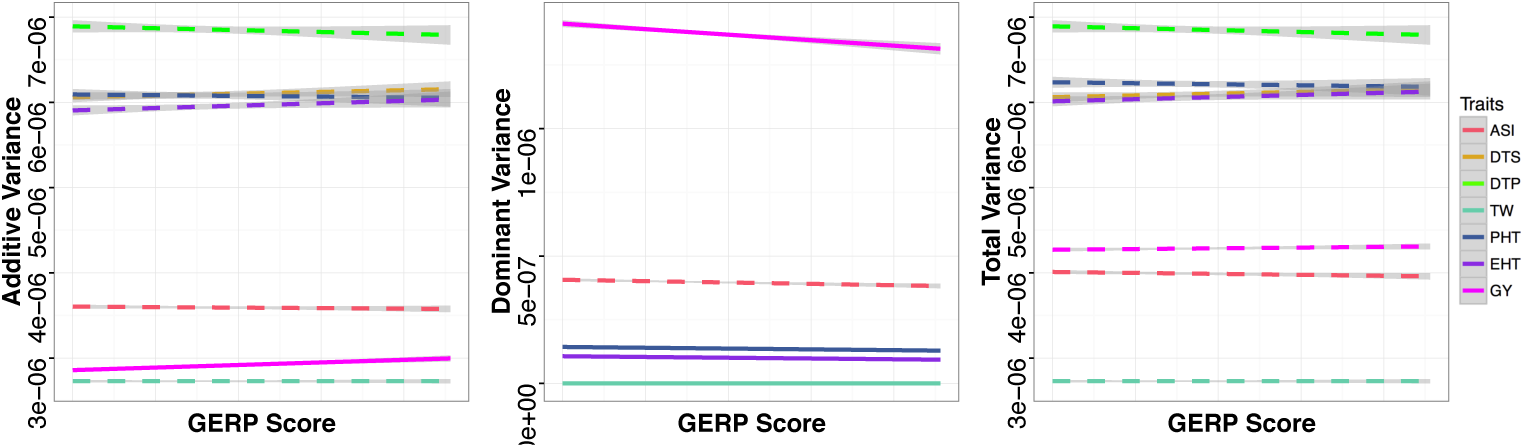
Linear regressions of GERP-SNPs’ additive variance, dominance variance and total variance of seven traits *per se* against their GERP scores. Solid and dashed lines represent significant and non-significant linear regressions, with grey bands representing 95% confidence intervals. Data are only shown for SNPs with > 1× of the mean genome-wide phenotypic variance explained.

**Fig S9.**
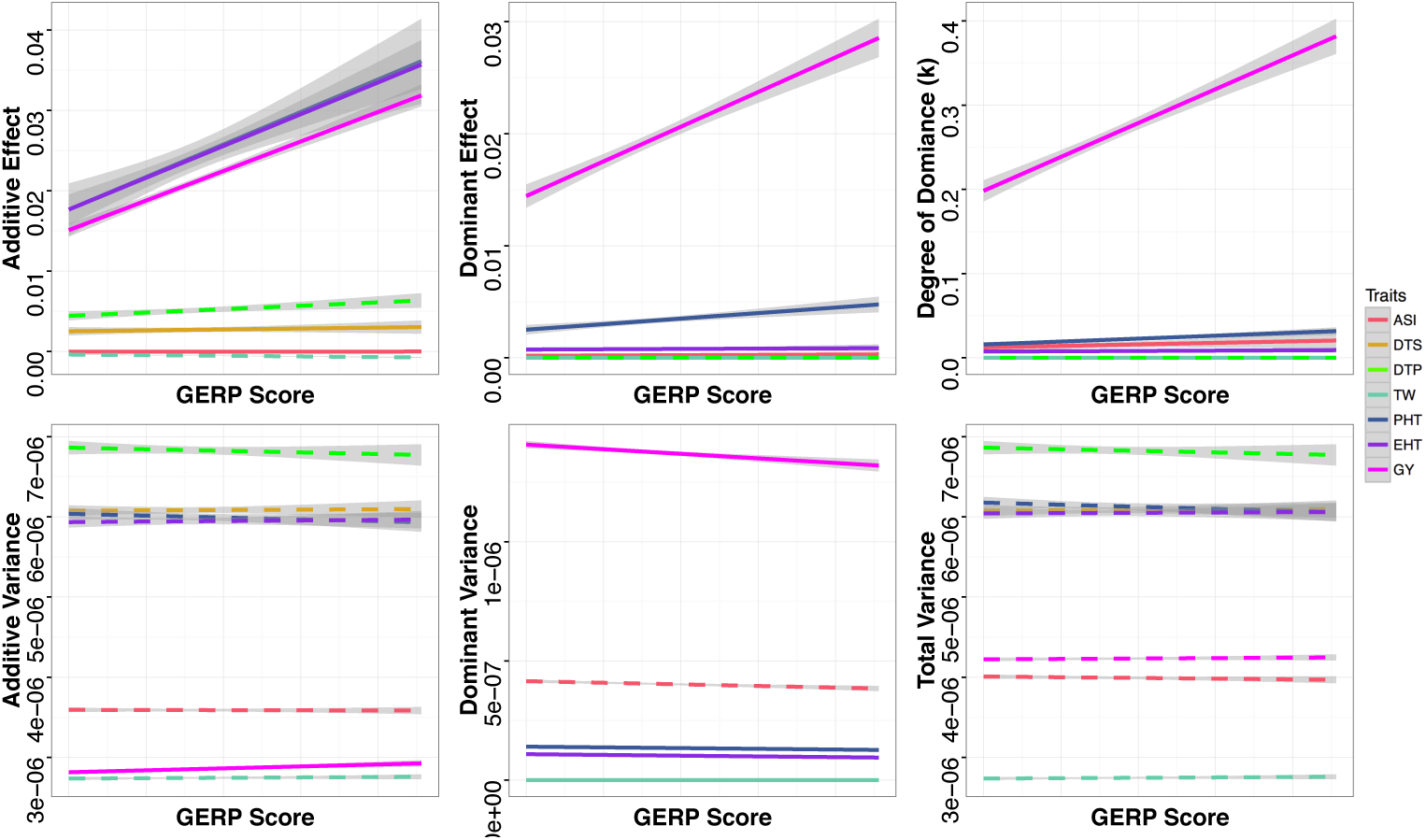
Linear regressions after filtering out GERP-SNPs located in regions in the lowest quartiles of recombination. Solid and dashed lines represent significant and non-significant linear regressions, with grey bands representing 95% confidence intervals. Data are only shown for GERP-SNPs with > 1× of the mean genome-wide variance explained and with > 1st quantile of the recombination rate (cM/Mb).

**Fig S10.**
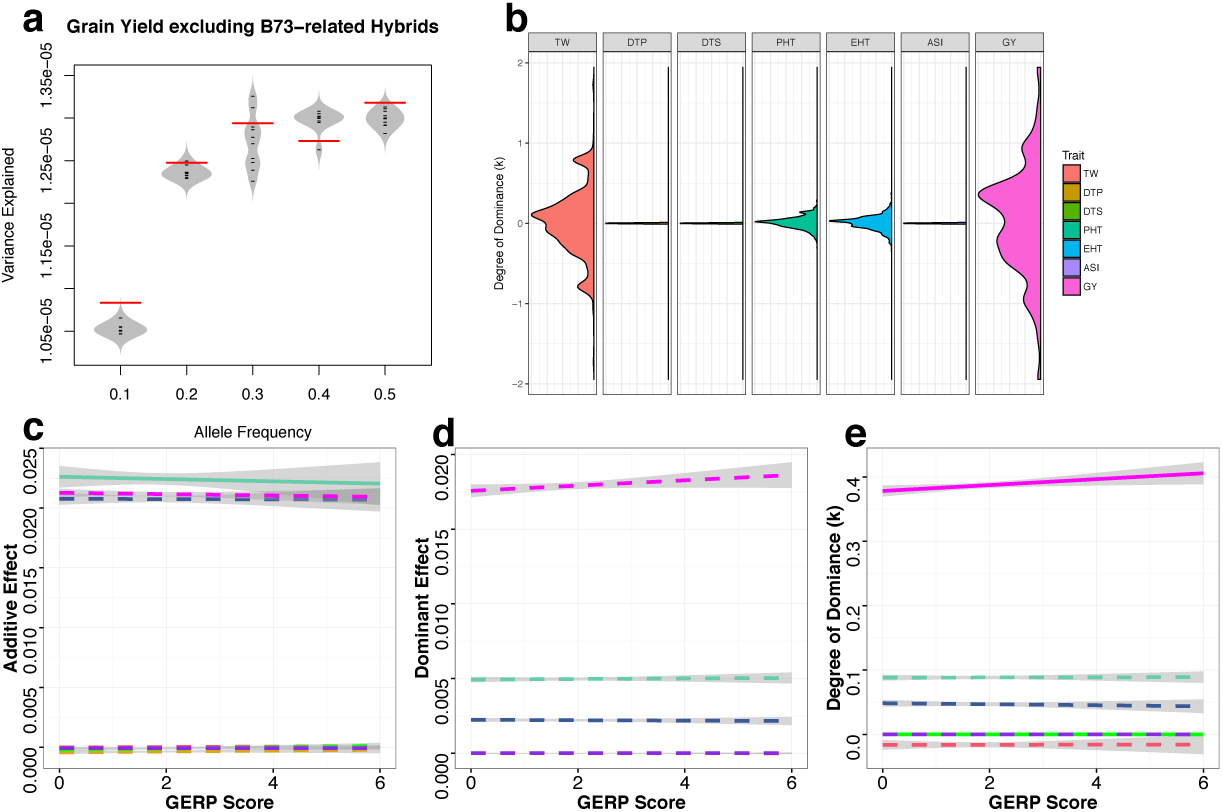
Phenotypic variance explained for grain yield and degree of dominance (*k*) of GERP-SNPs after removing 11 hybrids that B73 as one parent. **(a)** Total per-SNP variance explained for grain yield trait *per se* by deleterious (red lines) and randomly sampled SNPs (grey beanplots). **(b)** Density plots of the degree of dominance (*k*). Extreme values of *k* were truncated at 2 and −2 for visualization. **(c-e)** Linear regressions of additive effects **(c)**, dominance effects **(d)**, and degree of dominance **(e)** of seven traits *per se* against SNP GERP scores. Colors in **(c-e)** are the same as the legend for **(b)**. Solid and dashed lines represent significant and nonsignificant linear regressions, with grey bands representing 95% confidence intervals. Data are only shown for deleterious alleles with the above mean genome-wide variance explained.

**Fig S11.**
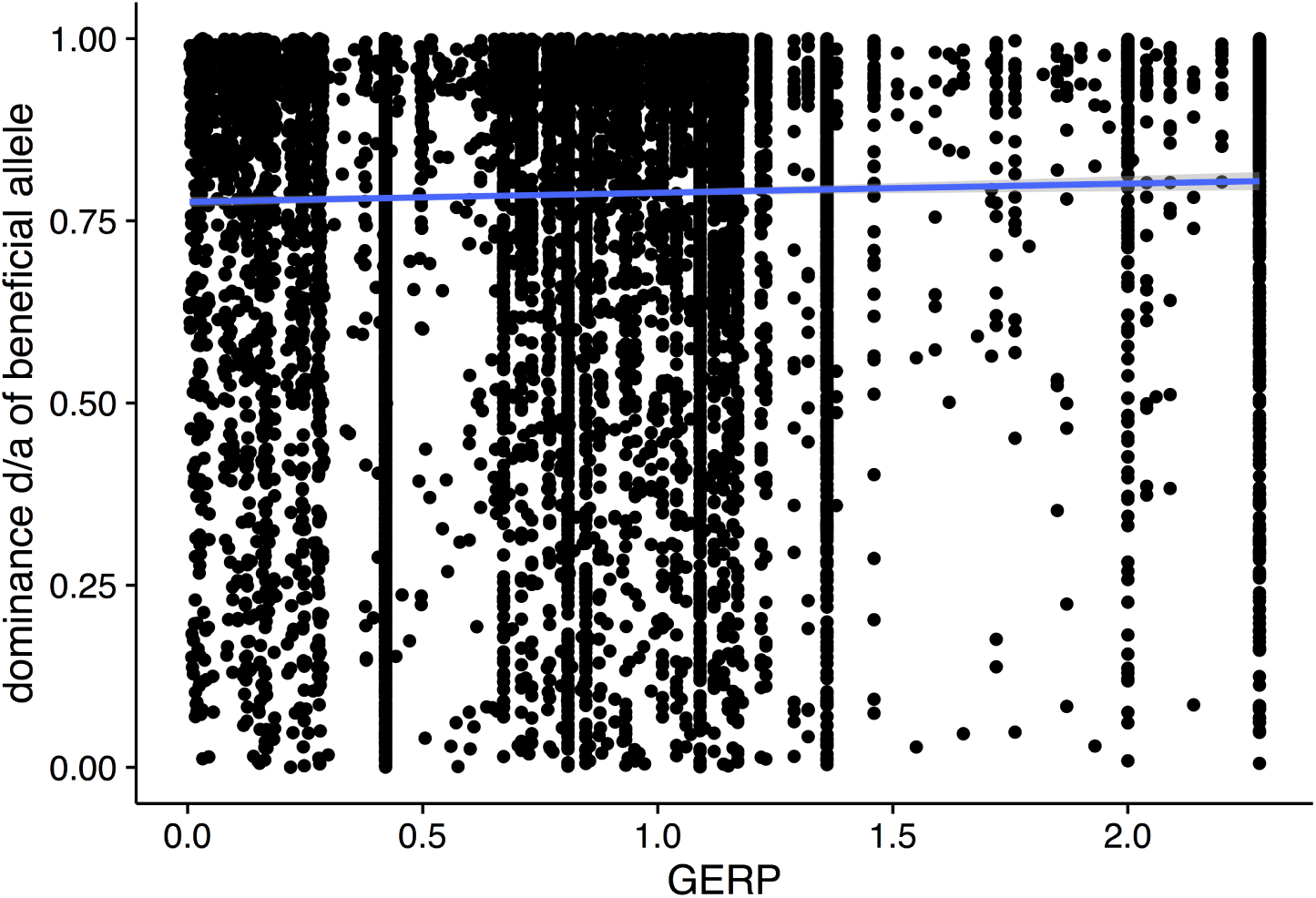
Regression of degree of dominance (*k*) on GERP scores for simulated data. Solid blue line indicates the regression line fitted to data simulated under mutation-selection balance (see Methods for details).

**Fig S12.**
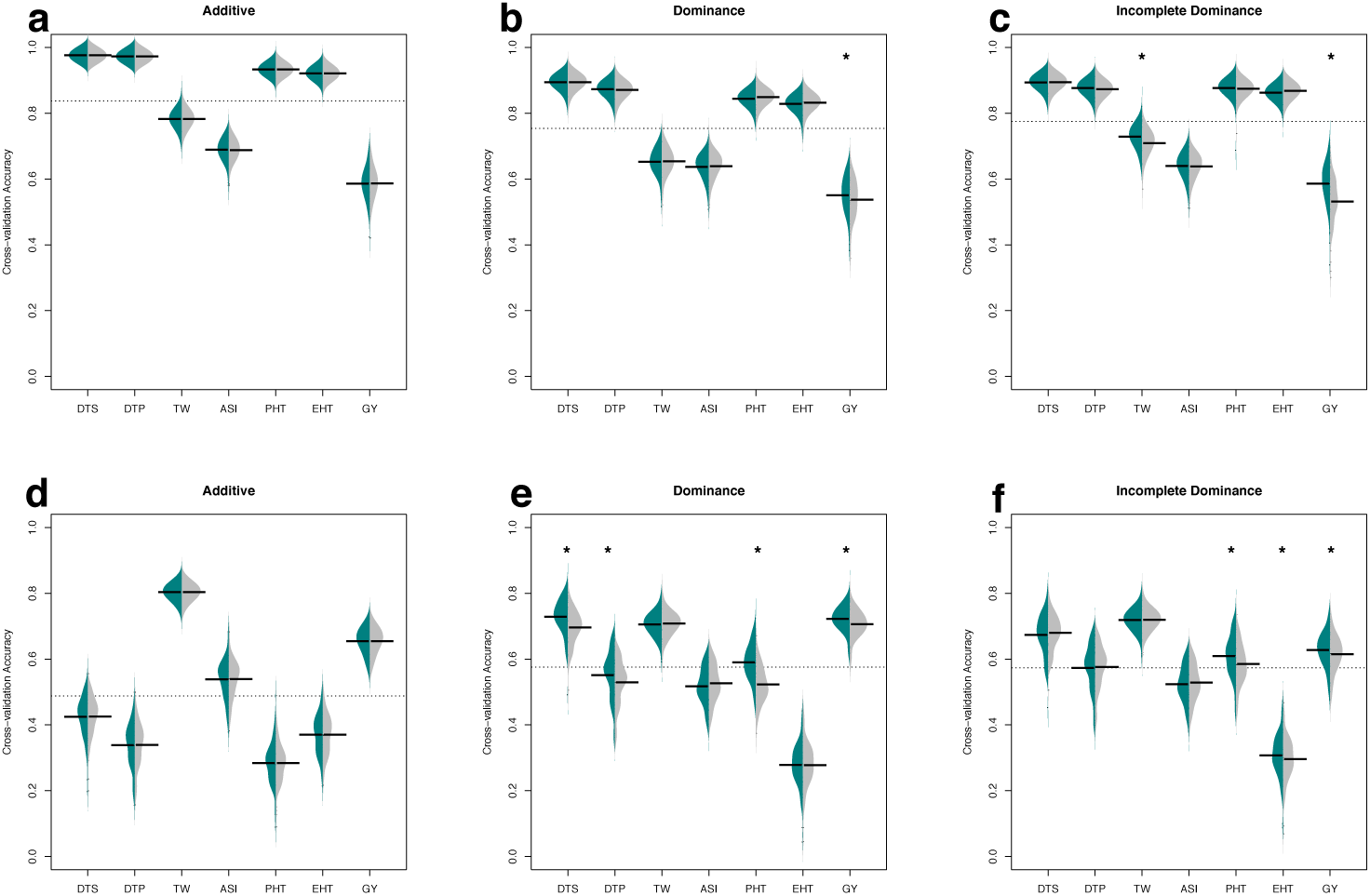
Cross-validation accuracy using GERP-SNPs in genic regions. Beanplots represent prediction accuracy estimated from cross-validation experiments for traits *per se* (**a, b, c**) and heterosis (**d, e, f**) under additive (**a, d**), dominance (**b, e**), and incomplete dominance (**c, f**) models. Prediction accuracy using real data is shown on the left (green) and permutation results on the right (grey). Horizontal bars indicate mean accuracy and the grey dashed lines indicate the overall mean accuracy. Stars indicate significantly (permutation FDR < 0.05) higher than cross-validation accuracy.

**Fig S13.**
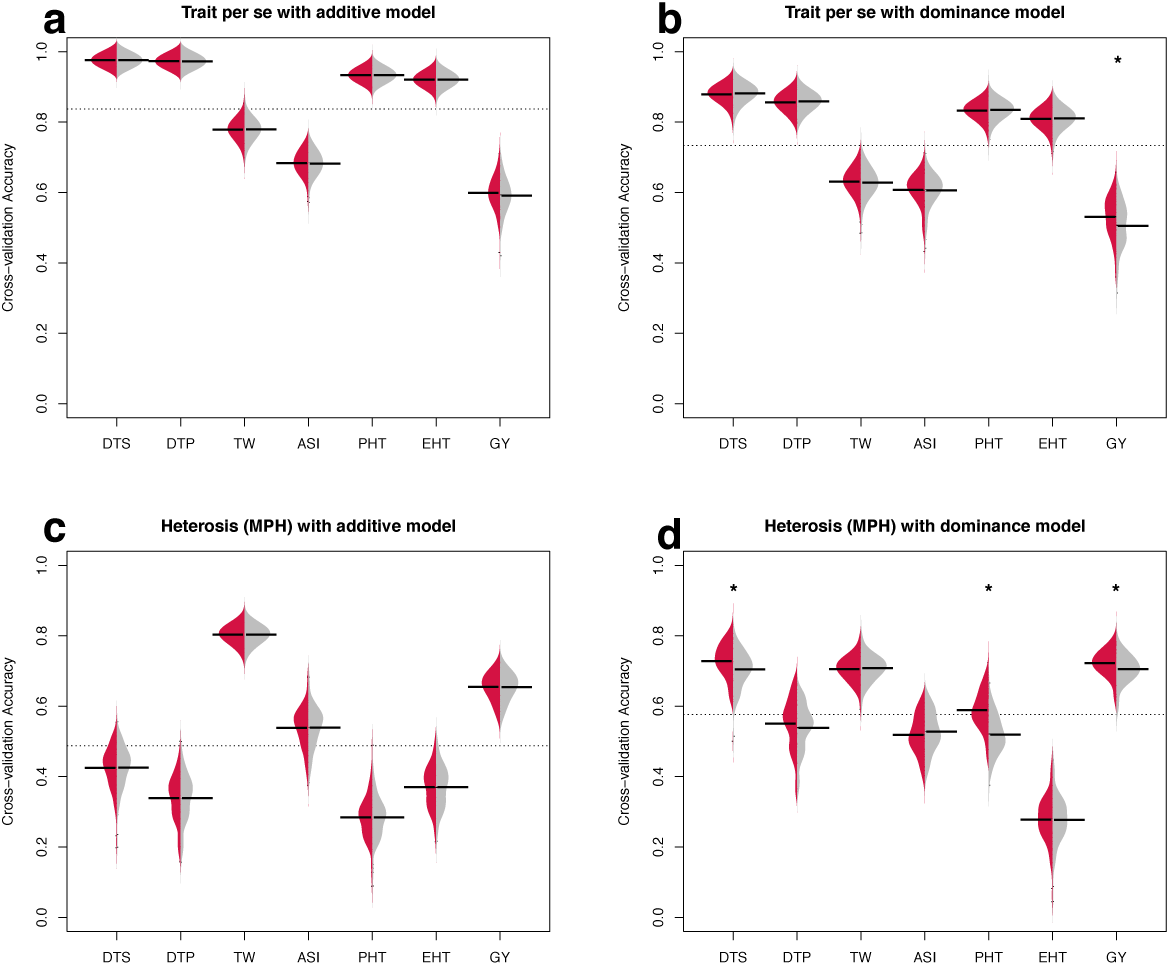
Cross-validation prediction accuracy for trait *per se* and heterosis. Beanplots represent prediction accuracy estimated from cross-validation experiments for traits *per se* (**a, b**) and heterosis (**c, d**) under additive (**a, c**) and dominance (**b, d**) models. Prediction accuracy using real data is shown on the left (red) and permutation results on the right (grey). Horizontal bars indicate mean accuracy of each trait and the grey dashed lines indicate the mean accuracy of all traits. Stars indicate real data having significantly (t-test P value < 0.05) higher cross-validation accuracy than permuted data.

**Fig S14.**
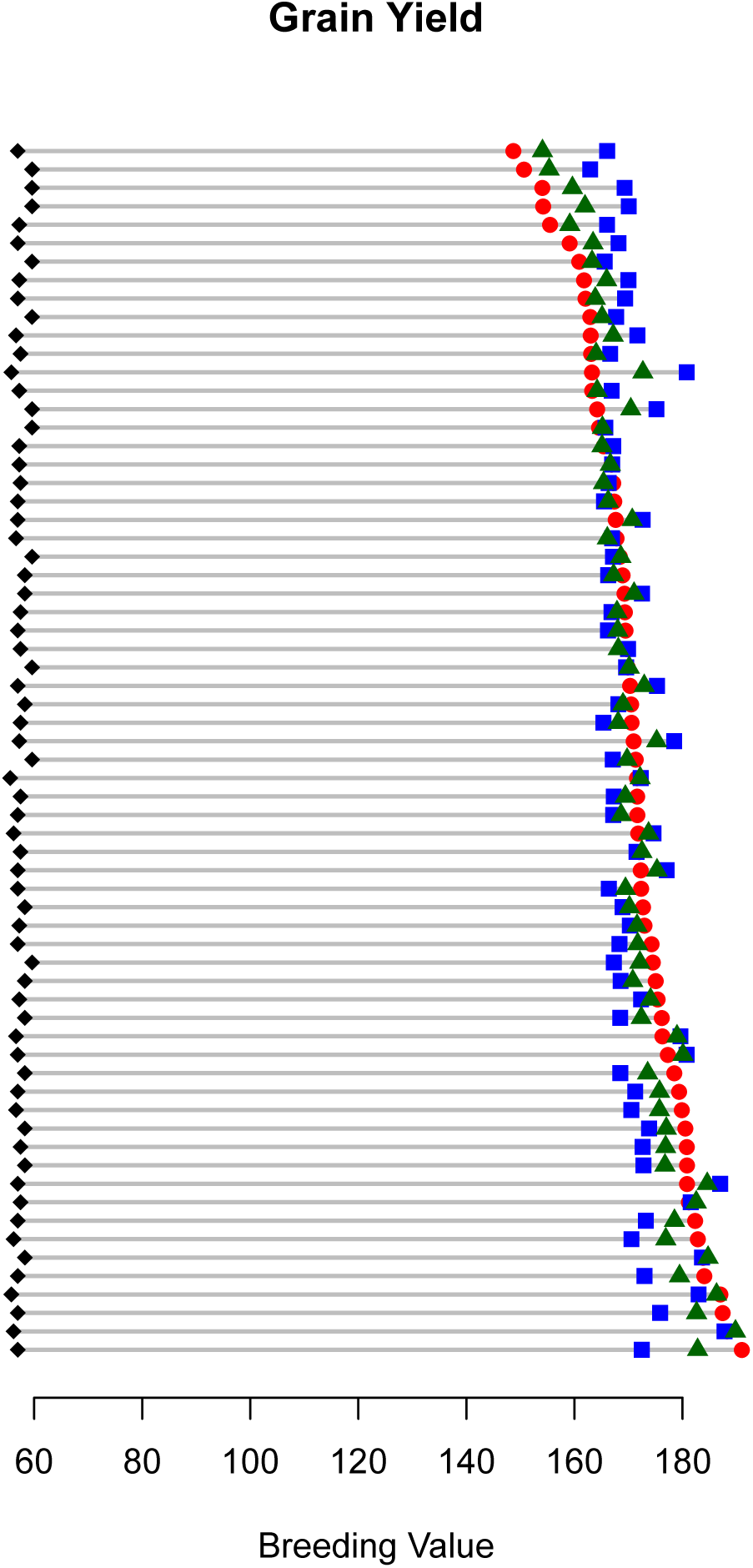
Breeding values of grain yield for diploid and simulated triploid hybrids. Each line represents the posterior breeding values of a diploid hybrid (red circle), its best parent (black diamond), and predicted breeding values of AAB triploid (blue square) and ABB triploid (green triangle) based on estimated effect sizes and dominance values for each SNP.

